# HIF1A modulate glycolysis function to governs mouse ovarian microenvironment metabolic plasticity in aging single cell resolution

**DOI:** 10.1101/2022.03.01.481557

**Authors:** Xiaoyu Zhang, Sumedha Gunewardena, Yan Liu, Ning Wang

## Abstract

The molecular machinery of ovarian aging and female age-related pathway remain unclear. Here, we utilized single-cell RNA-seq to profile over 9815 cells from both young and old female mouse and identified age-related alterations in the female somatic microenvironment. Interestingly, by aging-related signature calculation, we examined HIF1A in mouse ovarian cell aging regulated roles which effect pathways included glycolysis, TCA, OXPHOS and fatty acid metabolism. Additionally, inactivated HIF1A, decreased glycolysis was observed. Comparison analysis reveals the aging related regulon; metabolic and nutrient absorption changes provides a comprehensive understanding of the cell-type-specific mechanisms underlying mouse ovarian aging at single-cell resolution. This study, revealing new potential candidate biomarkers for the diagnosis of aging-associated ovary pathology.

## Main

Female reproductive system, the ovary, serves as a powerful model to study related between aging and metabolism^1^. Despite recent evidence that mammalian females supplied new oocytes during adult life, the role of ovarian aging is still problem which is poorly understood^2–4^. During parental age increased, aging-associated problems are related to multiple risk factors including reduced fertility and higher miscarriage rates^5^. Aging adversely affects genome integrity, epigenetic status and endocrine disorder in both male and female^6^. Thus, the ovary and testis are indispensable for the maintenance fertility and endocrine homeostasis^7–9^. An in-depth understanding of the mechanism, which drive ovarian aging is critical important. Although many studies have revealed aging-related alters in both male and female, it is still poorly unknown how aging impacts the mouse ovary at the molecular and genomic level by single cell resolution^10–12^.

Hypoxia inducible factor 1a (HIF1a), which belongs to the bHLH-PAS family of transcription factors, a factor adapts to hypoxic microenvironments functionally. However, it is puzzling that, although recent study demonstrated that HIF-1a activate gene pathways that minimize oxygen consumption, reduce reactive oxygen species (ROS), and restore oxygen delivery in heart, the role of HIF1a in ovary aging^13^. HIF1a is expressed in mouse ovary. Interestingly, this gene was observed impact preovulatory follicles function, inhibition of the HIF transcriptional activity blocked ovulation by preventing the rupture of the preovulatory follicles^14^. Now, much studies are focused on understanding the oxygen regulated function of HIF1a in the ovary. However, the other effect by HIF1a and the relationship between HIF1a and aging have not to be evident yet. In normal physiologically ovary, low oxygen concentration is beneficial for reproductive, cells prefer hypoxic environments and use glycolysis to produce ATP. Therefore, focusing on aging ovary metabolism, especially under low oxygen conditions, glycolysis may contribute to a good direction for study HIF1a role in aging ovary^15^.

In early studies, conventional bulk RNA-seq protocol have more flaws in accurately demonstrating changes from heterogeneous organ, such as ovary, in gene expression level and cell types alteration. By using advances single cell RNA sequencing (scRNA-seq) approach it is possible to accurately analyze alterations at the molecular and genomic level by single cell resolution within highly heterogeneous tissues. For our study, we used aging mouse combine with mouse organ scRNA-seq database to survey the first comprehensive single cell transcriptomic landscape of mouse ovarian aging. Overlap with aging-related signature calculation, we identified seven transcriptional factors in mouse ovarian cell aging regulated roles. Moreover, one of these aging-associated transcriptional factors changes (HIF1a) revealed that glycolysis was an essential factor of ovarian aging. To identified how HIF1a effect glycolysis, a multiple analysis with developmental mouse ovary RNA-seq data and a Hif1a knockout ChIP-seq and RNA-seq data revealed similar aging-associated downregulation of HIF1a and glycolysis genes. Our data provide potential biomarkers for the clinical diagnosis of ovarian aging. Single cell transcriptional profiling reveals a key role for the HIF1a-glycolysis axis in mediating ovarian aging, which provides a target for therapy aging-associated ovarian disorders and female infertility.

## Methods

### Animals

All animal experiments were conducted according to the approved protocol by the Institutional Animal Care and Use Committee at the University of Kansas Medical Center in strict accordance with its regulatory and ethical guidelines. All animals were housed in a specified pathogen-free facility with a 12□h light/dark cycle. All animals had access to food and water ad libitum. CD1 and DBA/2 mice were raised under specificpathogen-free (SPF) condition.

### RNA isolation and qPCR

Total RNA was isolated from collected cells using TRIzol^™^ (Invitrogen,15596018). The RT reaction was carried out with SuperScript II First-Strand Synthesis Kit (Invitrogen, 18080-051). qPCR was performed with gene specific primers that were listed in Supplementary Table 3. qPCR with amplified cDNAs was performed using the Power SYBR Green Master Mix (Applied Biosystems) on Applied Biosystems Quant Studio 5.

### Immunofluorescence (IF) analysis

For the IF staining, cultured cells were fixed with 4% paraformaldehyde contained 0.1% Triton X-100 for 10□min at room temperature. Then cells were washed with PBS. Blocking was performed using 5% BSA for 1□h at room temperature. The primary antibodies were added and incubated for overnight at 4□°C. After washed in PBS, the secondary antibodies were added and incubated for 1□h at room temperature. Images were captured using Nikon A1R confocal microscope and were processed using Nikon NIS Elements and Adobe Photoshop.

### Antibodies for immunofluorescence

Primary antibodies used were: GAPDH (CST, 5174) 1:200 dilution; HK2 (CST, 2867) 1:200 dilution; LDHA (CST, 3582) 1:200 dilution; PKM (CST, 3190) 1:200 dilution; PFKP (CST, 8164) 1:200 dilution; HIF1A (CST, 36169) 1:200 dilution; P53 (CST, 5174) 1:200 dilution. Secondary antibodies, donkey anti-mouse, donkey anti-rabbit, or donkey antigoat antibodies conjugated to AlexaFluor-488, AlexaFluor-546 or AlexaFluor-647, were purchased from Thermo Fisher and used at 1:500 dilution.

### Single-cell RNA-seq (scRNA-seq)

Treated cells were digested by collagenase IV for 20□min at room temperature. The digestion was then stopped by media. The cells were pelleted by centrifugation at 300□×□g for 5□min. And then, supernatant was removed, and the cell pellet was washed with PBS twice. Single cells were obtained by filtering through 40□μm strainers. Cell number was counted using Countess II FL automated cell counter (Invitrogen). The 10× Genomics Single-Cell 3’ Expression library preparation is performed using the 10× Genomics Chromium Controller. The cells prepared from disassociated tissue or tissue culture are validated for viability and cell concentration using the Countess II FL Automated Cell Counter (Life Technologies) targeting ≥75% cell viability. If debris or cell clumping is present in the cell suspension, the preparation is filtered through a FLOWMI Cell Strainer, 40□μm (Thermo Fisher 50-136-7555) to yield a homogenous single-cell suspension. Cell counts are re-determined by using the Countess ll FL and adjusted to ~1000□cells/μl by low speed centrifugation at 4□°C and re-suspended in 1× PBS without calcium or magnesium (Thermo Fisher MT21040CV) supplemented with 0.04% BSA to prepare cells for emulsification. The cell emulsification is performed with the 10× Chromium Controller using the Chromium Next GEM Single-Cell 3’ GEM Library & Gel Bead Kit v3.1 (10× Genomics 1000120) and Chromium Next GEM Chip G Single-Cell Kit (10× Genomics 1000127). Well 1 of Chip G is loaded with the RT Master Mix□+□cell suspension containing ~16,000 cells to target 10,000 emulsified cells at ~65% efficiency of emulsification. Well 2 of Chip G is filled with 50 μl of the Next GEM GEL Beads. Well 3 of Chip G is filled with 45□μl Partitioning Oil. Any unused wells are filled with 50% glycerol at a volume designated for the well number. A gasket is applied to the Chip G and the loading cassette and inserted into the Chromium Controller for GEM creation Using the Chromium Single-Cell G run program. Emulsified GEMs are recovered from each well and transferred to 200□μl strip tubes. The RT reaction to generate 10× barcoded single stranded cDNA, in the single-cell containing GEMs, is performed using an Eppendorf MasterCycler Pro thermal cycler. Post GEM-RT Cleanup is conducted by breaking the GEMS with 10X Recovery Agent to separate the aqueous phase from the Recovery Agent and Partitioning oil. A cleanup of the sscDNA containing aqueous phase is completed using Dynabeads MyOne Silane beads (Life Technologies 37002D). The second strand cDNA amplification is performed using the cDNA Amplification Mix on the Eppendorf MasterCycler Pro. The 3’ gene expression library construction is initiated with fragmentation, end repair and A-tailing of the dscDNA followed by adapter ligation and a sample index PCR using the Illumina compatible indexed adapters in the Chromium i7 Multiplex kit (10× Genomics 120262). Validation of the single-cell library is conducted using the Agilent Tapestation 4200 ScreenTape assay (Agilent 5067-5576). Single-cell library quantification is completed using a Roche LightCycle96 using FastStart Essential Green Master (Roche 06402712001) and KAPA Library Quant (Illumina) DNA Standards (KAPA KK4903). Library concentrations are adjusted to 3 nm and pooled. The library pool is diluted to 1.00□nm for a final clustering concentration of 200pM on a NovaSeq6000. The sequencing was performed using a NovaSeq6000 100 cycle Reagent Kit (Illumina 20012865) with an asymmetrical sequencing profile (read 1—28 cycle: i7 index read—8 cycle: i5 index read—0 cycle: read 2—94 cycles). Bcl2fastq conversion and demultiplexing is performed using the 10X Genomics Cell Ranger and Loupe Browser software suite.

### Analysis of single-cell RNA-seq data

Cell clustering was performed by Seurat (https://satijalab.org/seurat/, R package, v3.1). Seurat object was created first. Then, we discarded low-quality cells, in which less than 200 genes were detected. Genes expressed in less than three cells were filtered out. We also filtered cells that have lower than 2000 genes and that contain mitochondrial genome higher than 10% of mapped reads. We then used LogNormalize, a global-scaling normalization method, which normalized gene expression measurements by the total expression per cell, followed by multiplication of the result by a default scale factor (10,000) and subsequent log-transformation. After filtering and normalization, 9815 cells were selected for further analysis. Gene expression was calculated by centering expression across all cells in the cohort using the “ScaleData ()” function in Seurat. Identification of top 2000 variable genes, PCA^16^ (30 significant PCs determined by a scree plot) and SNN-Cliqinspired clustering were performed in Seurat to generate uniform manifold approximation and projection (UMAP) visualizations. Cluster definition was performed by specific marker genes and the “FindClusters ()” function in Seurat. Gene expression analysis for different clusters were performed by using the Monocle^17^ (http://cole-trapnell-lab.github.io/monocle-release/, R package, v2.12) under the default settings. For the gene expression analysis, we used a published code from a published study with modifications. Pheatmap^18^ package was used for heatmap plotting and clustering. GO analysis of genes were analyzed by using the ClusterProfiler^19^ package.

### Statistical analysis

All experiments were replicated at least three times independently. Different mice, tissues or cells were used during each experimental replicate. Quantitative data from the experimental replicates were pooled and are presented as the mean□±□SEM or mean□±□SD as indicated in the figure legend. Compiled data were analyzed by Student’s t test and two-way ANOVA test.

### Data availability

The authors declare that all data supporting the findings of this study are available within the article and its Supplementary Information files or from the corresponding author upon reasonable request. FastQ files of single-cell RNA-seq are available on Gene Expression Omnibus (GEO) database under accession code GSE???. Source data are provided with this paper.

### Code availability

Source code of the analysis is publicly available on GitHub at this address: https://github.com/iamzhangxiaoyu/scRNA-seq and are available on Zenodo: https://doi.org/10.5281/zenodo.4535405.

## Results

### Single Cell Transcriptome Identified Cell Type, Distribution and Gene Expression of the Aging Mouse Ovary

Characterizing the diversity of the ovary, we obtained aging mouse ovary tissues from 20 health female mouse of same age for 2 separated 10X genomic single cell RNA-seq assay. In addition, we integrated our aging mouse ovary 10X genomic data with published adult ovary data from mouse cell atlas^20^ by microwell-seq with the batch effect removed by harmony package^21^ in R. To further understand the aging and young mouse ovary, a total of 9815 cells were retained for further analysis after filtering by using stringent quality control (Extended Data Fig.1a). For the defining of each cell type, we processed the sequencing data by the DoubleFinder package^22^ firstly for the doublet remove and Seurat package^23^ for normalization, clustering and annotation of each cell type based on the expression of specific marker gene including granulosa cell (GC, *Inha*^+^), stroma cell (SC, *Col1a2*^+^), theca cell (TC, *Cyp11a1*^+^), ovary surface epithelium cell (OSE, *Krt19*^+^), immune cell (IC, *Lyz2*^+^) and endothelial cell (EC, *Cldn5*^+^) (Fig.1c,f). We have identified 6 major clusters, which visualized by uniform manifold approximation and projection (UMAP)^24^, and showed that the distribution in young and aging mouse ovary (Fig.1a, Extended Data Fig.1d,e). Histologically analysis (via PAS and Masson staining) revealed that thickness of the ovaries collected from aged mouse elevated compared to young (Fig.1d,e). To further investigate the proportion of cell, we calculated the average cell number and cell proportion in different cell types and stages (Fig.1b, Extended Data Fig.1b,c). Analysis of biological function of each cell cluster in mouse ovary surface by Gene Ontology (GO)^25^ using differentially expressed genes (DEGs) revealed the specific and enrichment function in different cell cluster (Extended Data Fig.2a,b,3a,b). For example, GO term enriched to SC include extracellular matrix organization and collagen fibril organization. GO term including T-cell activation and leukocyte proliferation were enriched for IC (Fig.1g). GSVA analysis^26^ showed that different score in hallmark function among the different cell clusters. Most of hallmark function score were consistent with the GO term enrichment. Such as DNA repair function score was the highest in GC which was also enriched in GO term (Fig.1h).

**Figure 1.**
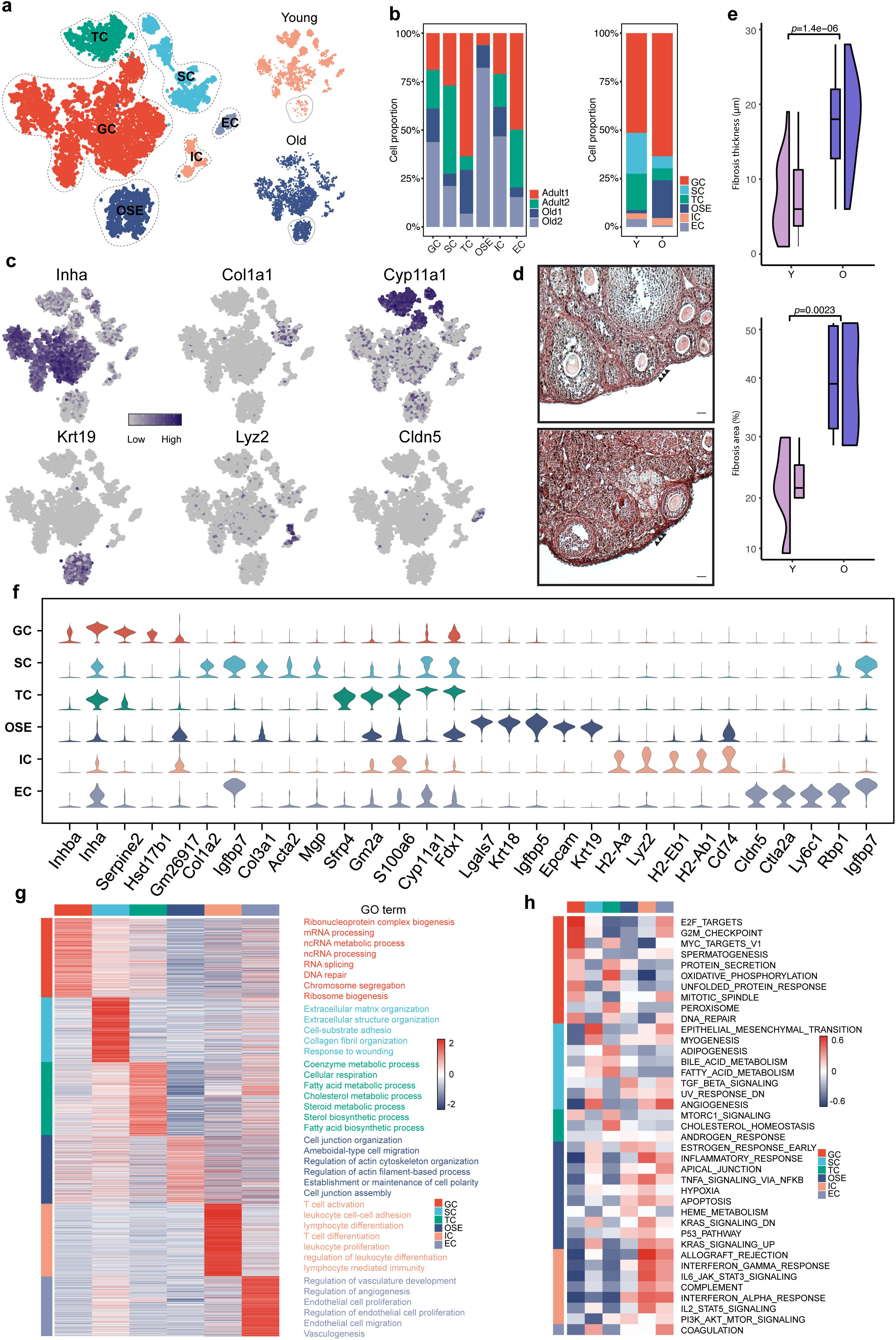
scRNA-seq analysis of young and old female ovary. (**a**) Uniform manifold approximation and projection (UMAP) plot of ovarian cells captured from young and old female ovary colored by cluster and sample identity. (**b**) Histograms showing the percentage of each sample and stages in each ovarian cell cluster. (**c**) Different stage germ cell marker expression shown as UMAP feature. (**d**) Picrosirius red (PSR) staining of mouse ovarian sections from young and old, respectively, demonstrating morphological changes during aging and variations among old mouse ovary. (**e**) PSR staining data are presented for the young and old group. The blue box indicates the old thickness and the fibrosis area, the purple box indicates the young thickness and the fibrosis area. (**f**) Violin plot showing the expression of representative genes for each cell type. (**g**) Heatmap showing gene expression signatures of each cell type. Enriched GO terms annotation for each cell type are on the right. (**h**) Heatmap showing hallmark GSVA analysis results of each cell type.

### DEGs and regulons in Aged Mouse Ovary

Initially, we identified aging-associated DEGs expression in mouse ovary surface in old versus young cell types GC, SC, TC, OSE, IC and EC, respectively (Fig.2a). Further analysis of DEGs expression, we calculated 387 downregulated and 1619 upregulated DEGs in Young vs Old group by function ‘FindAllMarkers’ (|avg-logFC|>0.25 and *p*_val_adj<0.05) in Seurat package (Fig.2b and Supplementary Table 1). Next, to study aging genes function in mouse ovary, we defined an aging-associated signature to distinguish young and old accounting for cell types from mouse ovary. To test enrichment of an aging-associated gene-set, we curated based on the hotspot genes which annotated in GenAge dataset of aging-related genes (Supplementary Table 2). 92 out of 388 aging-associated genes have been identified which represented in the single cell RNA-seq datasets to be aging associated genes. 85 out of 92 upregulated and 7 out of 92 downregulated genes in Young vs Old represented significant enriched upregulated and downregulated aging-associated genes in mouse ovary (Fig.2b). To summarized analysis these aging-associated genes in mouse ovary surface single cell RNA-seq datasets, we used ssGSEA^27^ (single sample Gene Set Enrichment Analysis) scores. After calculating both upregulated and downregulated aging scores, we showed broad variation from different cell types in mouse ovary (Fig.2c), suggesting that the variant in aging-associated genes that we observed is also cell type relevant. Transcription factors play critical roles in regulatory of gene expression. To identify regulons in young and old mouse ovarian cells, we used pySCENIC^28^ to calculate the regulatory activity of the GRNs. For characterizing of the regulatory patterns, we compared the difference of RAS scores of each regulon by the Connection Specificity Index^29^ (CSI) (Extended Data Fig.4a-f). Interesting, 209 regulons and 263 regulons are organized into 9 major modules, respectively in young and old mouse ovarian cells. (Fig.2d). For each module in young and old mouse ovary, we calculate the average activity scores by module and represented enriched cell types in modules. We found that some module specific enriched in cell types. In young ovary, module M1 and M2 contains regulons *Nr5a2*, *Gata6*, *Nfya* and *Klf5*, which are enriched in granulosa cells. M3, M4 and M7 contain regulons that are associated with metabolism and apoptosis, such as *Fos*, *Jun*, *Jund*, *Junb* and *Cebpb*, which are represented in ovary surface epithelial cells. Regulons *Clock*, *Nfyb*, *Stat3* and *Zbtb17* in M5 and M6 are associated with immune function enriched in immune cells. M8 and M9 are represented in endothelial cells and theca cells, respectively. In old ovary, regulons in M6 and M8 are highly associated with the nervous system. The activity of M6 and M8 including *Pparg*, *Six2*, *Foxo3* and *Rest* is specifically high in granulosa cells. Module M2 and M3 are related to ovary surface epithelial cells and they are very specifically enriched regulons *Erg*, *Rxra*, *Ddit3* and *Plag1*. Modules M1, M5 and M9 contains regulons that are activated in immune cells, such as *Elk1*, *Nrf1*, *Arnt*, *Hoxc10*, *Nfatc4* and *Pax8*. Regulons, *E2f1*, *Thap1*, *Nr1h4* and *Usf1*, are represented in theca cells (Extended Data Fig.5a,b). To further examine the difference in regulons between young ovary and old, we merged the regulons in young and old ovary. 73 out of 209 regulons enriched gene ontology (GO) functional terms and Kyoto encyclopedia of genes and genomes (KEGG) related to response to hypoxia and longevity regulating pathway in young ovary specifically. To compared with old ovary those regulons are 127 out of 263 which are associated with rhythmic process and cellular senescence (Fig.2e). A closer examination of these regulons with age associated DEGs, we represented a heatmap of 7 vital regulons in mouse ovary including *Cebpb*, *Myc*, *Trp53*, *Hif1a*, *Clock*, *Jund* and *Nfkb1*. We further list enriched DNA-binding motifs specific in mouse ovary surface (Fig.2f,g). For validation, we stained Hif1a, which are expressed by GC and EC. And, we found both its expression level and signal elevated in Young compare to Old. Number and fluorescence intensity of these two TFs were decreased in aged GCs in comparison with GCs from young ovaries (Fig.2h).

**Figure 2.**
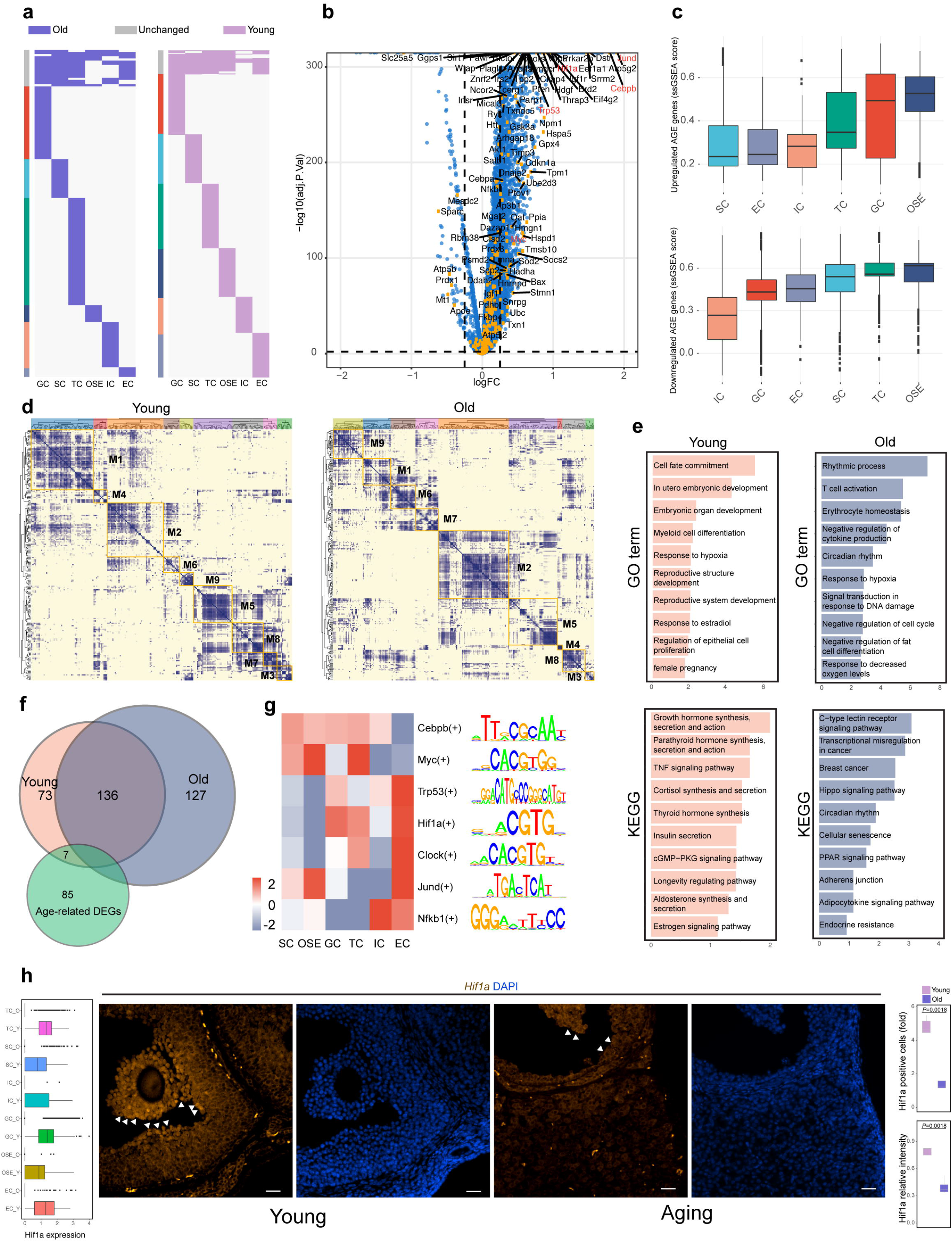
DEGs and regulons in young and aged mouse ovary. (**a**) Heatmaps showing common and unique upregulated and downregulated DEGs between young and old female ovary in each somatic niche cell type. (**b**) Volcano plot showing fold changes for genes differentially expressed between young and old pseudo-bulk samples. Age-related genes are enriched in upregulated and downregulated genes. (**c**) Boxplots of Aging-related-up and down enrichment scores show variation across ovarian somatic cell types. (**d**) Identified regulon modules based on regulon connection specificity index (CSI) matrix, along with representative transcription factors, corresponding binding motifs, and associated cell types in each young and old female mouse ovarian cells. (**e**) Top GO enrichments with representative young and old regulons. (**f**) Overlay of scRNA-seq regulons and aging related genes revealed 7 vital regulons in mouse ovary (**g**) Heatmap showing 7 vital regulons expression of each cell type. Binding site for each regulon are on the right. (**h**) Immunofluorescence analysis showing the downregulation of HIF1A in aged ovary in comparison to young counterparts. Scale bar, 50 μm. Box plots showing expression levels of *Hif1a* in young and old ovarian somatic cell. Quantification of HIF1A-positive cells and intensity were shown by box plots in mouse ovaries.

### Metabolic and Nutrient Absorptive Target Identification

To identified metabolic genes, we did pseudo-bulk analysis of single cell RNA-seq data by group young and old mouse ovarian cells. We performed the metabolic pathway genes were commonly activated in young mouse ovary, including glycolysis, TCA, OXPHOS, fatty acid metabolism, and others (Fig.3a). To further analysis metabolic pathway in mouse ovary, we defined 4 gene set expression involve in glycolysis (n=63), TCA (n=31), OXPHOS (n=108) and fatty acid metabolism (n=42) as score for glycolysis signature, TCA signature, OXPHOS signature and fatty acid metabolism signature, respectively. The distribution of these 4 metabolic signatures was represented in Fig.3b. Meanwhile, we found these metabolic signature genes show broad variation across different cell types in mouse ovary surface, glycolysis signature, TCA signature, OXPHOS signature and fatty acid metabolism signature scores were significantly elevated in GC, TC and OSE compare to other cell types (Fig.3c). Due to impaired in nutrient absorption lead to multiple diseases in aging have been reported^30,31^. To map the distinct expression patterns of genes associated with nutrient absorption in aging, we calculated the ssGSEA scores relate to major nutrient transporters genes range from glucose, amino acids, vitamins, lipid, ions and inorganic solutes. As Fig.3d,e shown, genes related to vitamin were enriched in the SC, and genes involved in inorganic solute and bile salt were significantly elevated in TC. Genes contained water, amino acid and glucose were highly expression in the EC. There were no significantly changes in the genes involved in organic solute and ion. To compare young and old mouse ovary nutrient patterns, a dot plot was represented that genes related to nutrient absorption were significantly elevated in young mouse ovary which consistent with the expression patterns in metabolic pathway genes. Although there was no significantly fluctuation in the expression of organic solute and ion, some genes showed highly expression pattern in the young TC and OSE such as *Slc31a1* and *Kctd14*. In general, our data indicated that both metabolic pathway and nutrient absorption functional activated in young mouse ovary surface compare with old mouse.

**Figure 3.**
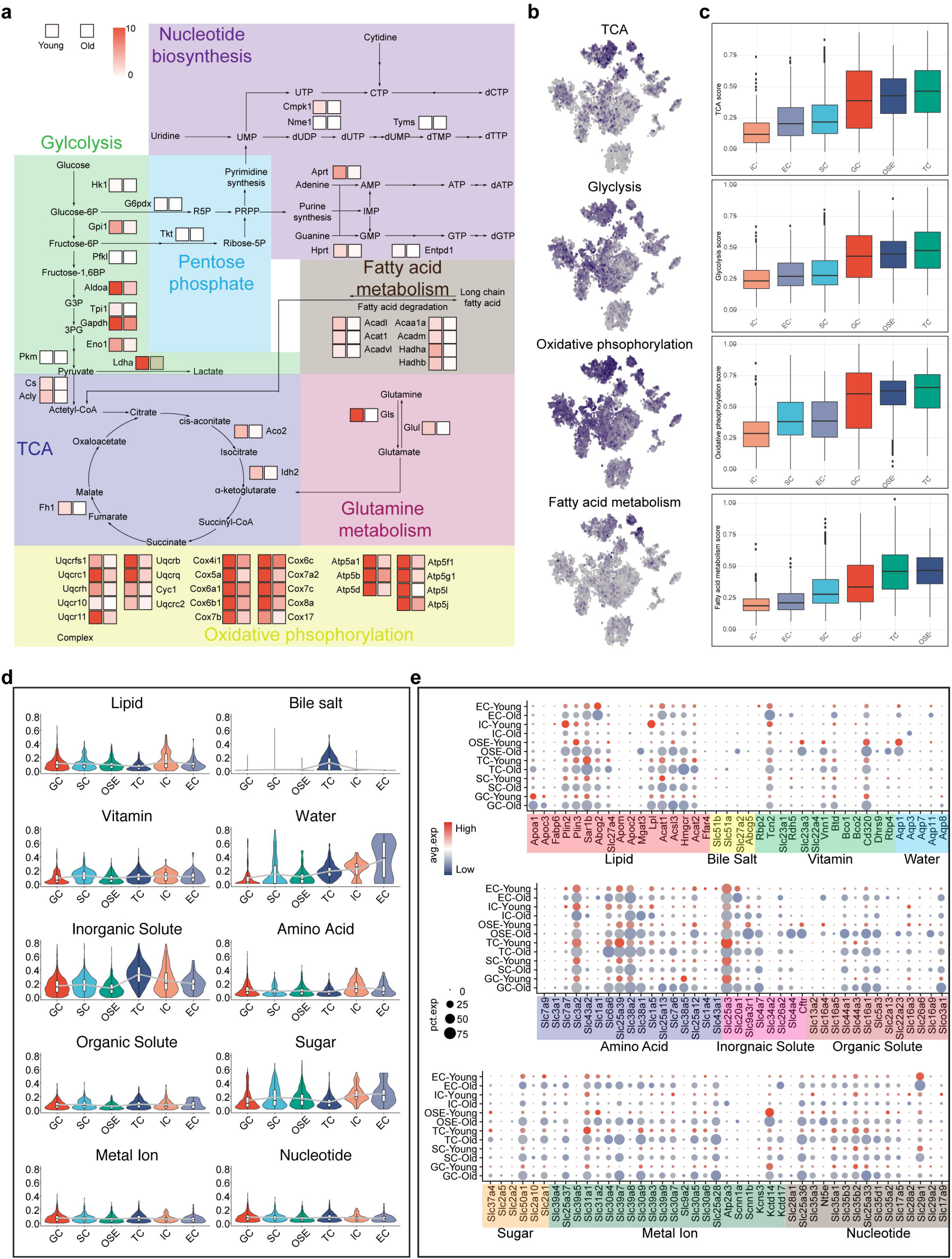
Metabolic and nutrient absorptive target identification. (**a**) Map of upregulated central carbon metabolic pathways in young versus old mouse ovary. Red color indicates upregulated expression. (**b**) Different metabolic signature marker expression shown as UMAP feature. (**c**) Boxplots of metabolic signature scores show variation across ovarian somatic cell types. (**d**) Violin plots showing distributions of mean expression of transporter genes in each cell type in mouse ovary (**e**) Expression patterns of specific genes involved in nutrient absorption and transport in each cell type in mouse ovary. Each dot represents a gene, of which the color saturation indicates the average expression level, and the size indicates the percentage of cells expressing the gene.

### Metabolic profiling identifies the role of glycolysis in Aged Mouse Ovary

Metabolic and nutrient absorptive target analysis revealed the differential function in young and aged ovaries, including glycolysis, TCA, OXPHOS and fatty acid metabolism. We next sought to identify glycolysis-associated changes in protein expression in ovarian cells. Glycolysis are essential for ovarian development and homeostasis provide nutrients and mechanical support for oocytes via physical interactions^32^. Due to glycolysis-related apoptosis of GCs often causes follicular atresia and ovarian aging^33^, we focused on the glycolysis-associated changes of gene expression in ovarian. Expression and protein levels of these glycolysis-associated genes were also decreased in aging ovarian cells in comparison with ovarian cells from young ovaries (Fig.4a-e). Immunostaining confirmed a significant reduction of HK2, LDHA, PKM, PFKP and GAPDH positive cells detected in young ovaries. qRT-PCR analysis showed a significant reduction of glycolysis-associated genes in in aging ovary compare young group (Fig.4f). Combined analysis of scRNA-seq with gene expression revealed glycolysis-associated genes decreased during ovary aging. Consistent with this observation of staining, the data suggest that glycolysis function may influence development in ovary function with aging.

**Figure 4.**
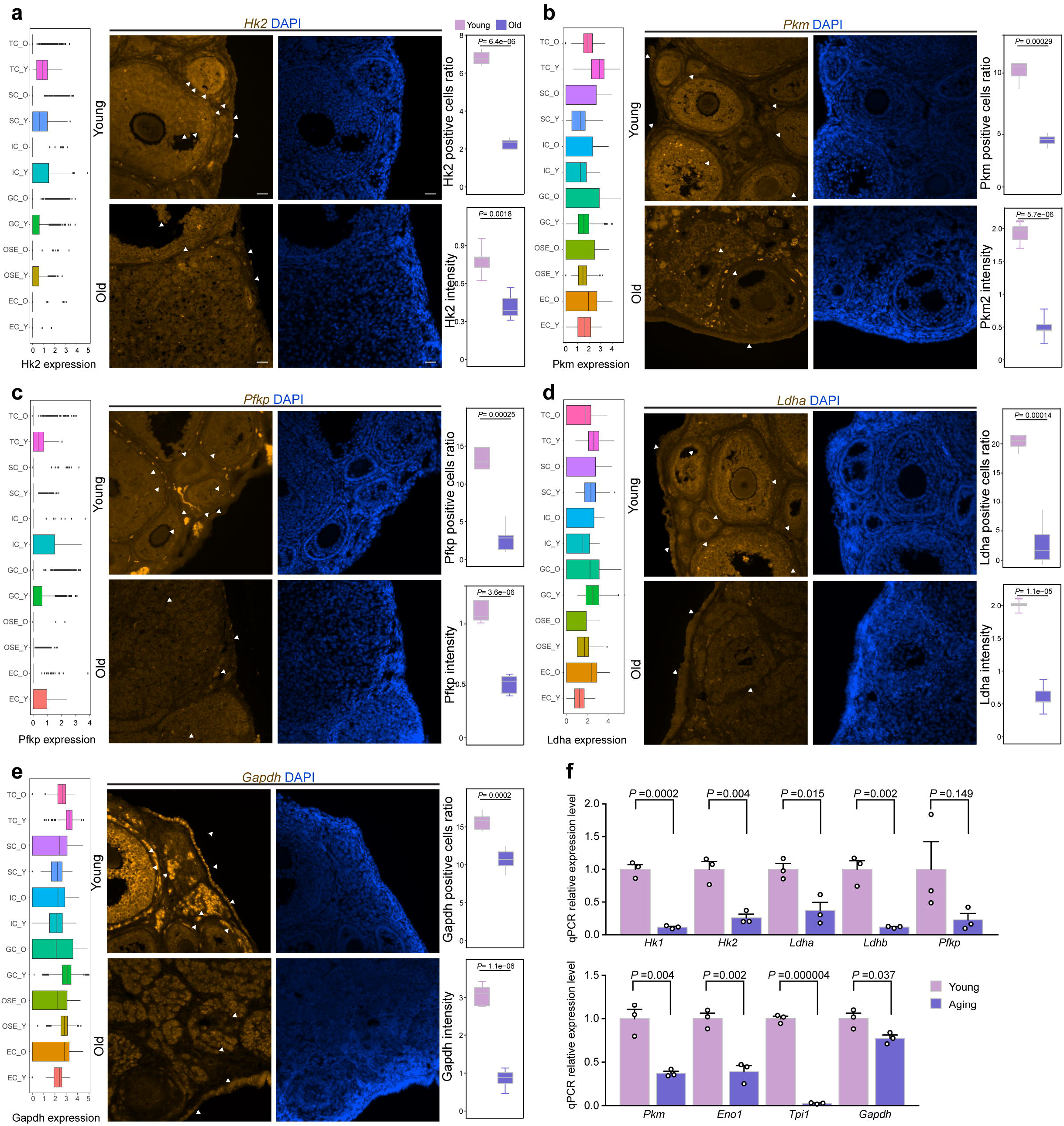
Metabolic profiling identifies the role of glycolysis in aged mouse ovary. (**a**) Immunofluorescence analysis showing the downregulation of HK2 in aged ovary in comparison to young counterparts. Scale bar, 50 μm. Box plots showing expression levels of *Hk2* in young and old ovarian somatic cell. Quantification of HK2-positive cells and intensity were shown by box plots in mouse ovaries. (**b**) Immunofluorescence analysis showing the downregulation of PKM in aged ovary in comparison to young counterparts. Scale bar, 50 μm. Box plots showing expression levels of *Pkm* in young and old ovarian somatic cell. Quantification of PKM-positive cells and intensity were shown by box plots in mouse ovaries. (**c**) Immunofluorescence analysis showing the downregulation of PFKP in aged ovary in comparison to young counterparts. Scale bar, 50 μm. Box plots showing expression levels of *Pfkp* in young and old ovarian somatic cell. Quantification of PFKP-positive cells and intensity were shown by box plots in mouse ovaries. (**d**) Immunofluorescence analysis showing the downregulation of LDHA in aged ovary in comparison to young counterparts. Scale bar, 50 μm. Box plots showing expression levels of *Ldha* in young and old ovarian somatic cell. Quantification of LDHA-positive cells and intensity were shown by box plots in mouse ovaries. (**e**) Immunofluorescence analysis showing the downregulation of GAPDH in aged ovary in comparison to young counterparts. Scale bar, 50 μm. Box plots showing expression levels of *Gapdh* in young and old ovarian somatic cell. Quantification of GAPDH-positive cells and intensity were shown by box plots in mouse ovaries. (**f**) qRT-PCR analysis of glycolysis related genes in young and old mouse ovary normalized to ß-actin. Data represent mean ± SD; n = 3 independent ovaries. P < 0.05 (two-way ANOVA test). Data are representative of three independent experiments.

### HIF1a signaling controls glycolysis

To determine whether HIF1a signaling controls glycolysis in ovary function with aging, we combined analysis transcript profiles of 3-mo, 6-mo, 9-mo, and 12-mo ovary with Hif1a knockout RNA-seq and ChIP-seq data. Initiation of the ovarian transcriptomic profile analysis from 3-month to 12-month age groups, RNA sequencing data were reanalysis (n =5/group. A heatmap shown of samples using all expressed genes demonstrated a separation of 3-mo, 6-mo, 9-mo and 12-mo ovary samples (Extended Data Fig.6a). All expressed genes were shown in three clusters through a K-mean clustering and a separation of 3-mo and 6-mo samples from 9-mo and 12-mo samples indicating a shift in transcriptomic patterns between young and aging ovary samples with both increases and decreases in expression observed. A line plot was demonstrated steadily linear increases or decreases with growing age across the 4 timepoints (Extended Data Fig.6b). To further annotated this set of genes, GO term function analysis was performed to identify Glycolysis, TCA, Oxidative phosphorylation and HIF-1 signaling pathway affected with aging. Among these pathways and processes were decreased broadly (Extended Data Fig.6c). Consistent with scRNA-seq, downregulated regulons found in aging (Hif1a, Trp53 and Cebpb) were shift down to the bottom by peak plot using RNA-seq data cross 3-mo to 12-mo ovary samples (Extended Data Fig.6d). To further identified the relationship between hif1a and glycolysis pathway, we analysis Hif1a knockout RNA-seq and ChIP-seq data^13^. Hif1a knockout RNA-seq show that Hif1a knockout exhibited comparable transcriptomic changes including shut down of essential glycolysis genes, such as *Gapdh*, *Ldha*, *Pfkp*, *Pkm* and *Hk2*, in response to Hif1a regulation (Fig.5a). GSVA analysis demonstrated that metabolic pathway concludes Glycolysis and Oxidative phosphorylation decreased in HIF1a knockout samples from GO term, KEGG, Hallmark and Reactome enrichment (Fig.5b). We also performed Gene Set Enrichment Analysis (GSEA) and found significant enrichment of genes associated with HIF-1signaling pathway and glycolysis in hif1a knockout mouse dataset (Fig.5c). Meanwhile we examined the region around *Hk2*, *Ldha*, *Eno1*, *Gapdh*, *Pkm* and *Pfkl* in Hif1a knockout RNA-seq and ChIP-seq data combine with growing aging ovary RNA-seq data (Fig.5d,e, Extended Data Fig.7a-c)^34^. Interestingly, the promoter specific bound peak demonstrated a strong binding region by Hif1a. Consistently, by examine Hif1a knockout RNA-seq data, we observed *Hk2*, *Ldha*, *Eno1*, *Gapdh*, *Pkm* and *Pfkl* expression level decreased at thses locus in Hif1a knockout samples. We also detected *Hk2*, *Ldha*, *Eno1*, *Gapdh*, *Pkm* and *Pfkl* gene expression decreased as well in aging ovary samples. This is consistent with these finding indicated that HIF1a signaling controls glycolysis in aging ovary.

**Figure 5.**
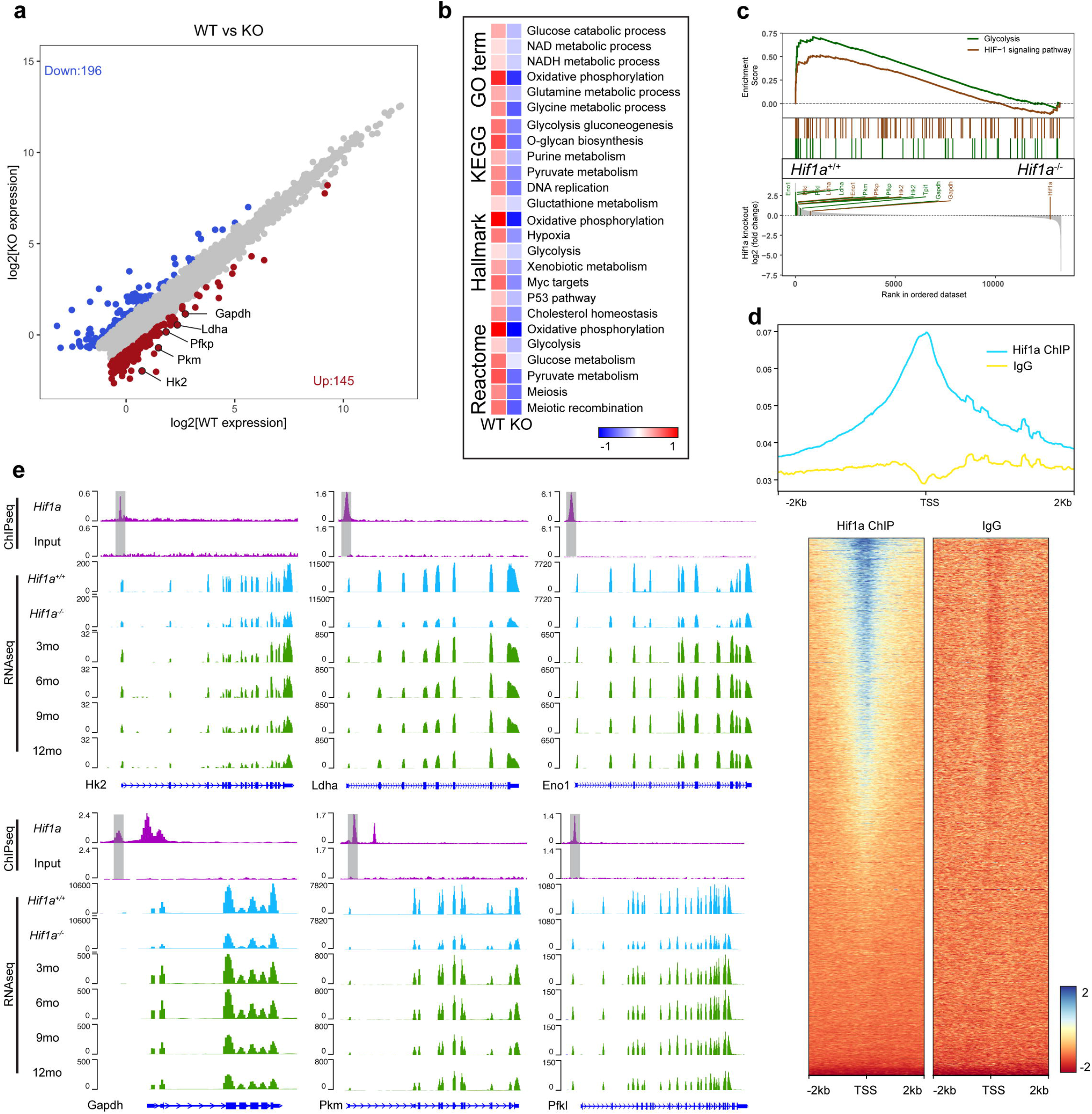
HIF1a signaling controls glycolysis. (**a**) Scatter plot representations of DEGs between Hif1a wildtype and Hif1a knockout. The up or down-regulated genes (fold change > 2 or < −2) in each group are plotted in red and blue, respectively. (**b**) GSVA analyses reveal distinct enriched gene sets between Hif1a wildtype and Hif1a knockout. In the heatmap, rows are defined by the selected gene sets, and columns by consensus scores for each group. (**c**) Gene Set Enrichment Analysis (GSEA) of the Hif1a wildtype and Hif1a knockout RNA-seq data. (**d**) ChIP-seq analysis of HIF1A bound. (**e**) Hk2, Ldha, Eno1, Gapdh, Pkm and Pfkl genome accessibility tracks for ChIP-seq and RNA-seq data.

### HIF1A interacts with glycolysis pathway

In order to identify essential proteins of the HIF1A interaction with glycolysis, we ranked proteins by their average functional similarity relationships among proteins within the interactome^35^. PFKL, PFKP and PKM were the three top-ranked proteins potentially playing central roles in the HIF1A interactome. PFKL, PFKP, PKM, TRP53, HK2 and LDHA protein with a cutoff value > 0.6 (Fig.6a). We further identified high confidence interactions (annotated by Kyoto Encyclopedia of Genes and Genomes (KEGG) or validated by low-throughput assays) more efficiently than co-expression based method. Interestingly, three glycolysis-associated interactors identified (PKM, HK2 and LDHA) were supported by KEGG or low-throughput experiments (Fig.6b). Then, we investigated HIF1A correlation with glycolysis pathway and HIF1A regulated glycolysis gene expression in the Genotype-Tissue Expression database (GTEx) (Fig.6c). Using HK2, ENO1, GAPDH, LDHA, PKM and PFKP as six representative glycolysis pathway genes, we found that HIF1A show strong (Pearson correlation > 0.6) or moderate correlation (Pearson correlation□>□0.4) with glycolysis pathway genes (Fig.6d). We further confirmed that HIF1A show high correlation with glycolysis pathway in our ovarian single cell RNA-seq dataset as well (Fig.6e).

**Figure 6.**
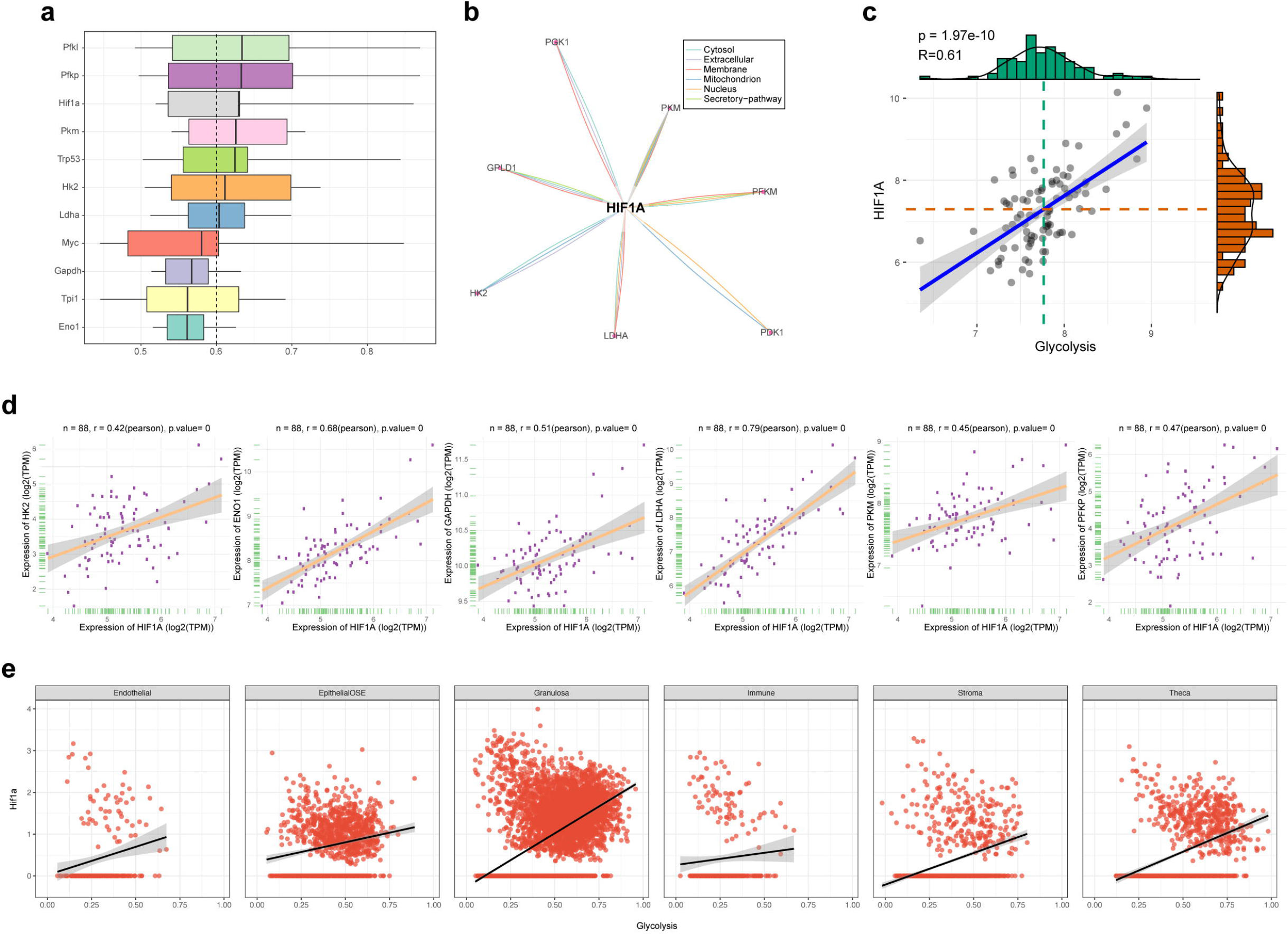
HIF1A interacts with glycolysis pathway. (**a**) The distributions of glycolysis related similarities were summarized as boxplots. The boxes represent the middle 50 percent of the similarities. Proteins with a higher average functional similarity (cutoff > 0.6) were defined as party proteins, which are considered as the central proteins within the HIF1A interactome. (**b**) The dashed line represents the cutoff value. The interactions predicted with high-confident interactions. (**c**) GTEx database showing correlation of representative HIF1A with glycolysis pathway. (**d**) GTEx database showing correlation of representative HIF1A with selected glycolysis related genes (HK2, ENO1, GAPDH, LDHA, PKM and PFKP) in 88 human ovary samples. The linear regression curve is demonstrated. 95% confidence interval (shaded area) is shown in each panel. Two tailed t-statistic P value and coefficient (R) of Pearson’s correlation is shown on the top. (**e**) scRNA-seq database showing correlation of representative HIF1A with glycolysis pathway in each cell type in female mouse ovary.

## Discussion

Due to fertility rates decline in aging mammalian, the cellular and molecular mechanisms are still challenging in ovarian aging. To understand the cellular and molecular mechanisms associated with ovarian aging, we representative scRNA-seq profiling of aging mouse ovary samples combine with young mouse ovary organ scRNA-seq database. In addition, we show similar genetic profiles in developmental mouse ovary RNA-seq data and a Hif1a knockout ChIP-seq and RNA-seq data which providing insights into the mechanisms. Our findings on alterations in ovary during aging are consistent with previous studies on developmental aging mouse ovary. Moreover, our study provided deep through scRNA-seq on the mouse ovary revealing key point of the HIF1a-glycolysis axis alterations at the molecular and genomic level in highly heterogeneous ovarian aging. Specifically, the downregulation of transcriptional factor Hif1a and downregulation of glycolysis pathways related to ovarian aging were two novel characters of aging ovary. Therefore, these observations provide novel insights in ovarian aging and identify target for the diagnosis and therapy of aging-associated ovarian disorders and female infertility.

Considering the cellular composited complication of the mouse ovary, we identified six ovarian somatic compartments based on their unique molecular marker. Previous studies have representative possible mechanisms aging-associated transcriptional changes, which largely involved the disturbance of antioxidant signaling and the induction of oxidative damage as a crucial factor in ovarian functional reduction with aging^36^. However, how ovarian functional decline and aging synergist effect on ovarian fertility is largely unknown, especially at the single cell level with such complicated heterogenetic composition. Here, we representative that each type of the soma compartment has extremely different aging related transcriptional switch relay on their specific aging related regulon, that are associated with ovarian functional impairment. To confirm these specific regulon, we show that ovarian metabolic function is linked to the specific cell-type-related downregulation of age-related regulons and metabolic signature. Correlation of metabolic changes with specific age-related regulons confirmed the robustness of the scRNA-seq strategy and reliability of our data. Consistently, bulk RNA-seq has revealed age-related metabolic dysfunction in mouse ovarian cells collected from 3mo to 12mo^34^.

According to scRNA-seq analysis, aging ovarian cells congruently downregulated the expression of metabolic marker genes, as well as metabolic genes involved in OXPHOS, glycolysis, TCA cycle, nucleotide biosynthesis, and mRNA metabolism. Hence, metabolic related cells seem to similarity enriched in GCs, OSEs and TCs. Nonetheless, tissue-specific differences in metabolic transcriptome profiles were observed, likely in part due to differences in micro-environmental conditions^37^. Our data also unveiled the differential expression of nutrient absorption–related genes in aging ovary across different cell types. High expression of the genes related to transport of lipid and sugar in the GCs and OSEs indicates that the absorption and application of these nutrients is mainly accomplished in GCs and OSEs, which is consistent with an earlier report^38^. Based on our scRNA-seq analysis, we representative that in mouse, the inactivation of the glycolysis occurs in aged ovarian somatic cells by 7 different regulatory mechanisms, evidenced by the transcriptional regulatory regulons involving aging related gene set. In a meanwhile, our single-cell transcriptomic atlas of mouse ovarian aging was mapped on samples collected from mouse ovaries with a wider sample numbers (n=20) which undoubtedly provides invaluable in-depth information to ovarian aging biology. Furthermore, the expression levels of glycolysis genes were positively correlated with aging related regulons in mouse ovarian cells, and Hif1a knockout responses metabolic change, highlighting Hif1a as biomarkers or targets for diagnose aging in clinic.

In summary, we profiled single cell transcriptomes of mouse ovarian cells from young and aging female mouse, which serves as a foundational dataset for the scientific community. Further comparison analysis reveals the aging related aging regulons, metabolic and nutrient absorption changes taking place during ovary aging, and provides potential candidate biomarkers for the diagnosis of aging-associated ovary pathology. In addition, the mechanistic insights arising from this study could establish new avenues for developing targeted metabolic interventions to protect against physiological ovarian aging and related diseases and for developing new guidelines to practice better lifestyle to improve fertility.

## Acknowledgments

Acknowledgments follow the references and notes list but are not numbered. Start with text that acknowledges non-author contributions and then complete each of the sections below as separate paragraphs.

## Funding

National Institutes of Health grant R01 HD103888 (NW)

KUMC Lied Pre-Clinical Research Pilot Grant Program (NW)

National Institutes of Health grant P20 GM103418 K-INBRE Postdoctoral Fellowship Award (XZ)

KUMC BRTP Postdoctoral Fellowship Award (XZ)

Department of Molecular and Integrative Physiology at KUMC (NW)

## Author contributions

Conceptualization: XZ, NW

Methodology: XZ, NW

Investigation: XZ, SG, YL

Visualization: XZ, NW

Funding acquisition: NW

Supervision: NW

Writing – original draft: XZ, NW

Writing – review & editing: XZ, NW

## Data and materials availability

All data, code, and materials used in the analysis must be available in some form to any researcher for purposes of reproducing or extending the analysis. Include a note explaining any restrictions on materials, such as materials transfer agreements (MTAs). Note accession numbers to any data relating to the paper and deposited in a public database; include a brief description of the data set or model with the number. If all data are in the paper and supplementary materials, include the sentence “All data are available in the main text or the supplementary materials.”

---

**Extended Data Fig. 1** 10X genomics quality control in scRNA-seq. (**a**) A workflow showing relationship between HIF1A with glycolysis. (**b,c**) Histograms showing the aveage number of each sample and stages in each ovarian cell cluster. (**d,e**) Uniform manifold approximation and projection (UMAP) plot of ovarian cells captured from young and old female ovary colored by assay method and sample identity.

**Extended Data Fig. 2** GO function and KEGG analysis for each cell type in scRNA-seq. (**a**) The bubble plots show GO biological process enrichment analysis for each cell type. (**b**) The bubble plots show GO molecular function enrichment analysis for each cell type.

**Extended Data Fig. 3** GO function and KEGG analysis for each cell type in scRNA-seq. (**a**) The bubble plots show GO cellular component enrichment analysis for each cell type. (**b**) The bubble plots show KEGG enrichment analysis for each cell type.

**Extended Data Fig. 4** Cell-Type-Specific Regulon Activity Analysis. (**a**) Rank for regulons in granulosa cell based on regulon specificity score (RSS). Granulosa cells are highlighted in the UMAP (red dots). Binarized regulon activity scores (RAS) for top rank regulon Foxo1 on UMAP (dark green dots). (**b**) Rank for regulons in endothelial cell based on regulon specificity score (RSS). Endothelial cells are highlighted in the UMAP (red dots). Binarized regulon activity scores (RAS) for top rank regulon Sox18 on UMAP (dark green dots). (**c**) Rank for regulons in OSE cell based on regulon specificity score (RSS). OSE cells are highlighted in the UMAP (red dots). Binarized regulon activity scores (RAS) for top rank regulon Grhl2 on UMAP (dark green dots). (**d**) Rank for regulons in theca cell based on regulon specificity score (RSS). Theca cells are highlighted in the UMAP (red dots). Binarized regulon activity scores (RAS) for top rank regulon Rora on UMAP (dark green dots). (**e**) Rank for regulons in immune cell based on regulon specificity score (RSS). Immune cells are highlighted in the UMAP (red dots). Binarized regulon activity scores (RAS) for top rank regulon Tbx21 on UMAP (dark green dots). (**f**) Rank for regulons in stroma cell based on regulon specificity score (RSS). Stroma cells are highlighted in the UMAP (red dots). Binarized regulon activity scores (RAS) for top rank regulon Prdm6 on UMAP (dark green dots).

**Extended Data Fig. 5** Activation of regulon modules in differnet cell types from young and old female mouse ovary. (**a**) Average module activity score in each cell type from young mouse ovary and enriched regulon binding sites. (**b**) Average module activity score in each cell type from old mouse ovary and enriched regulon binding sites.

**Extended Data Fig. 6** Bulk RNA-seq analysis of aging ovary. (**a**) A heatmap shown of samples using all expressed genes demonstrated a separation of 3-mo, 6-mo, 9-mo and 12-mo ovary. (**b**) A line plot was demonstrated steadily linear increases or decreases with growing age across the 4 timepoints. (**c**) GO term function analysis was performed to identify Glycolysis, TCA, Oxidative phosphorylation and HIF-1 signaling pathway affected with aging. (**d**) Peak plot using RNA-seq data cross 3-mo to 12-mo ovary samples.

**Extended Data Fig. 7** Hif1a ChIP-seq analysis. (**a**) Global analysis of distribution of Hif1a ChIP-seq binding site. (**b**) Global analysis of genome location of Hif1a ChIP-seq binding site. (**c**) Level of peak frequency in genomic region.

