## Supplementary figures and images for "HIF1A modulate glycolysis function to governs mouse ovarian microenvironment metabolic plasticity in aging single cell resolution"

### FigS1

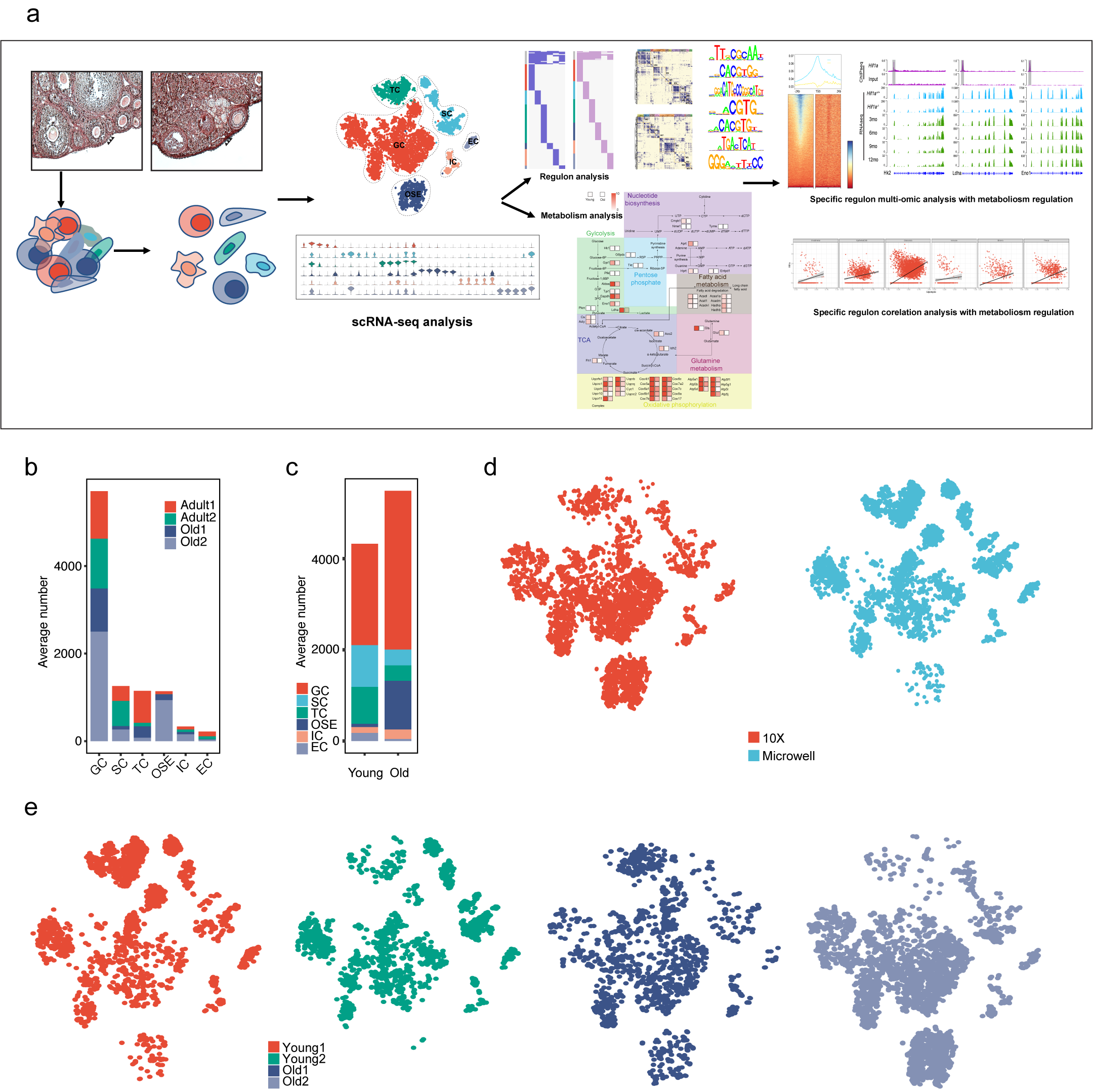

### FigS2

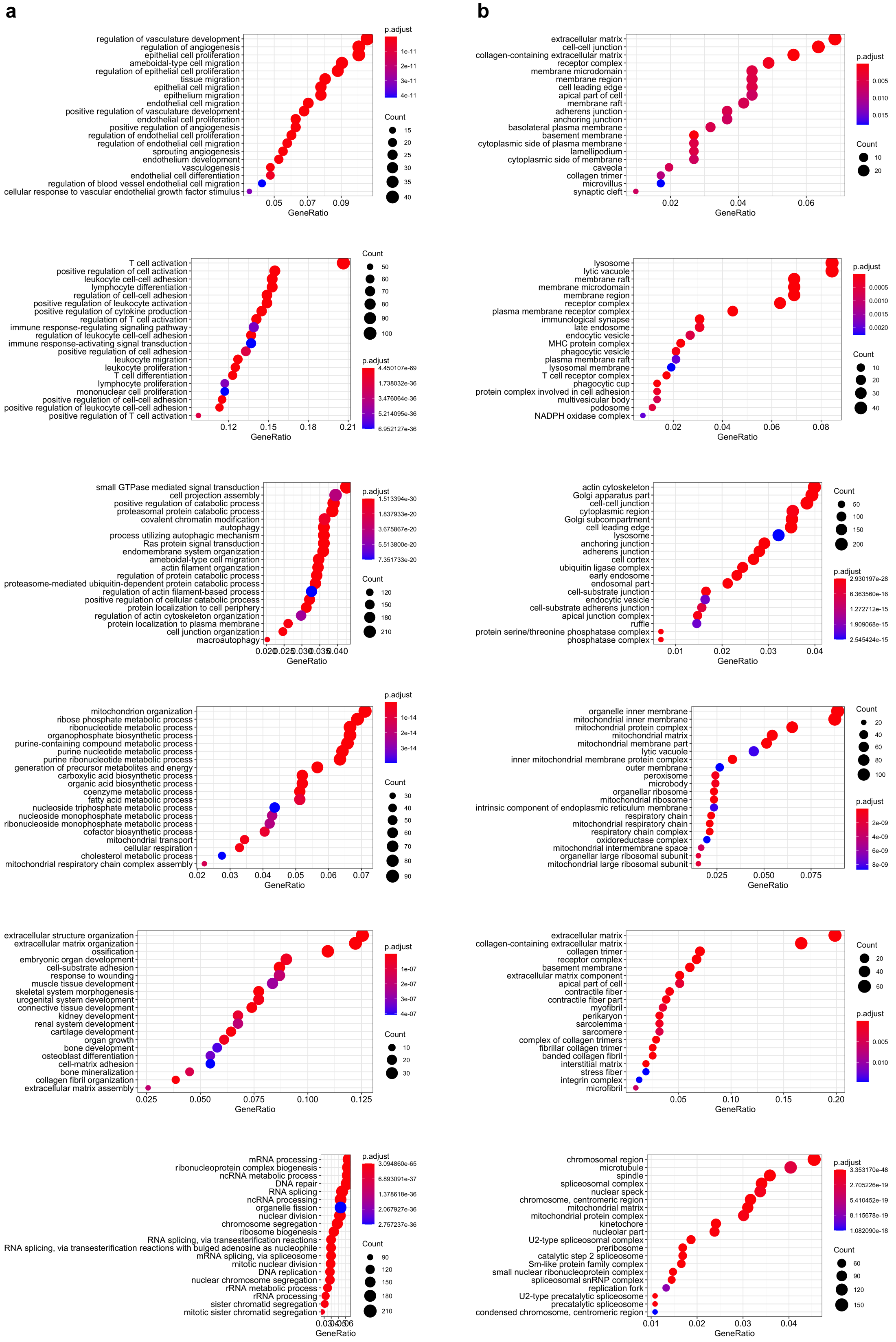

### FigS3

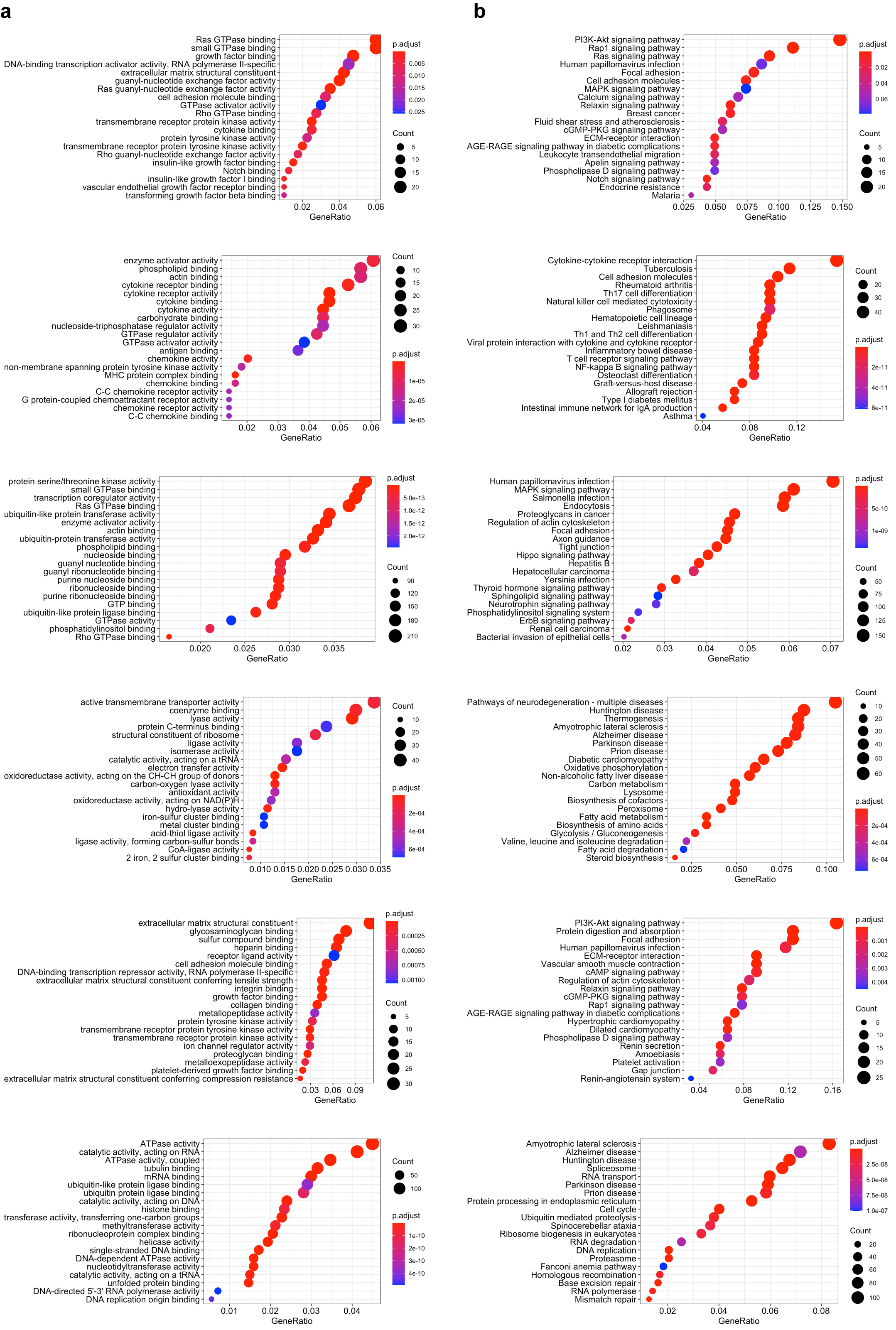

### FigS4

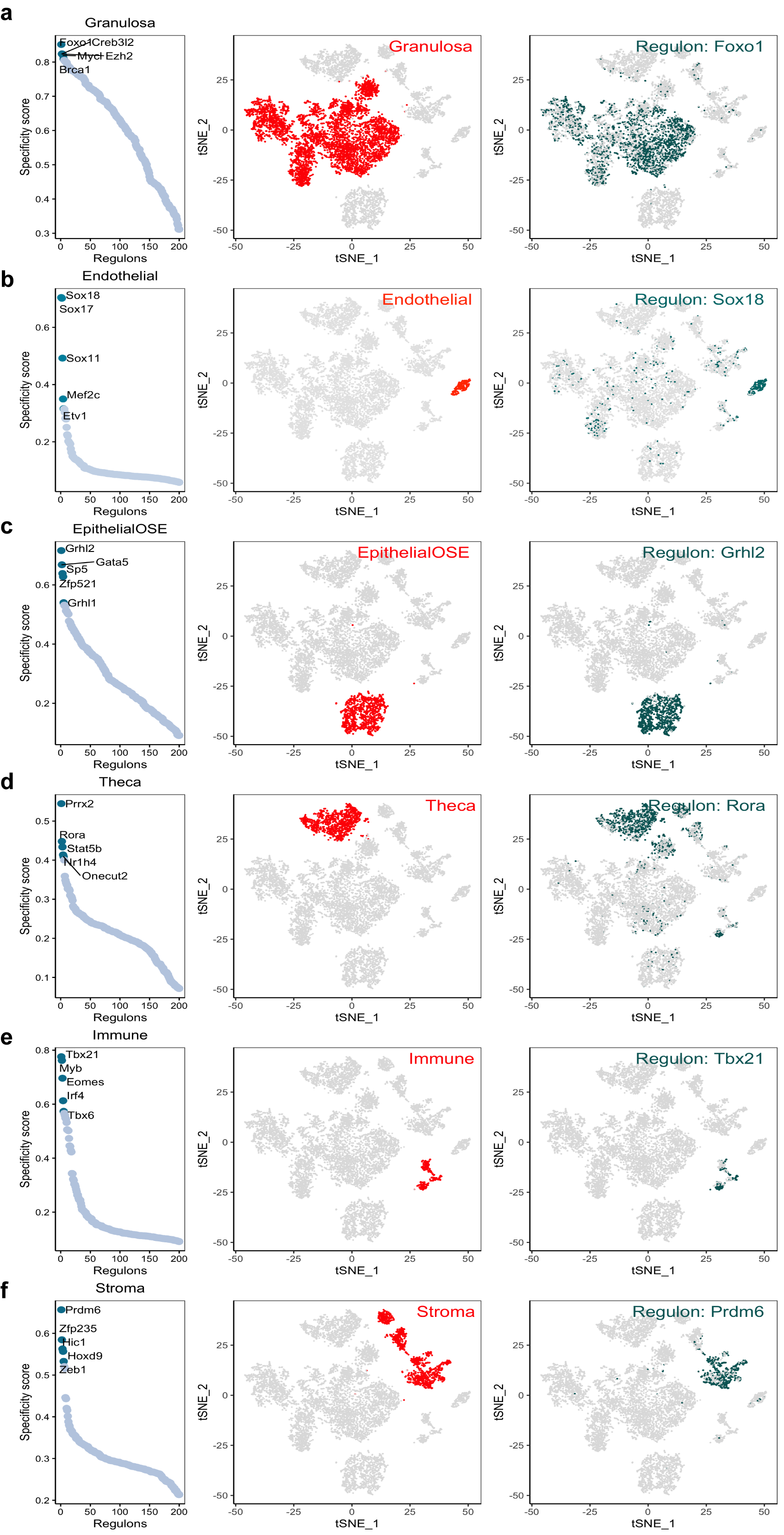

### FigS5

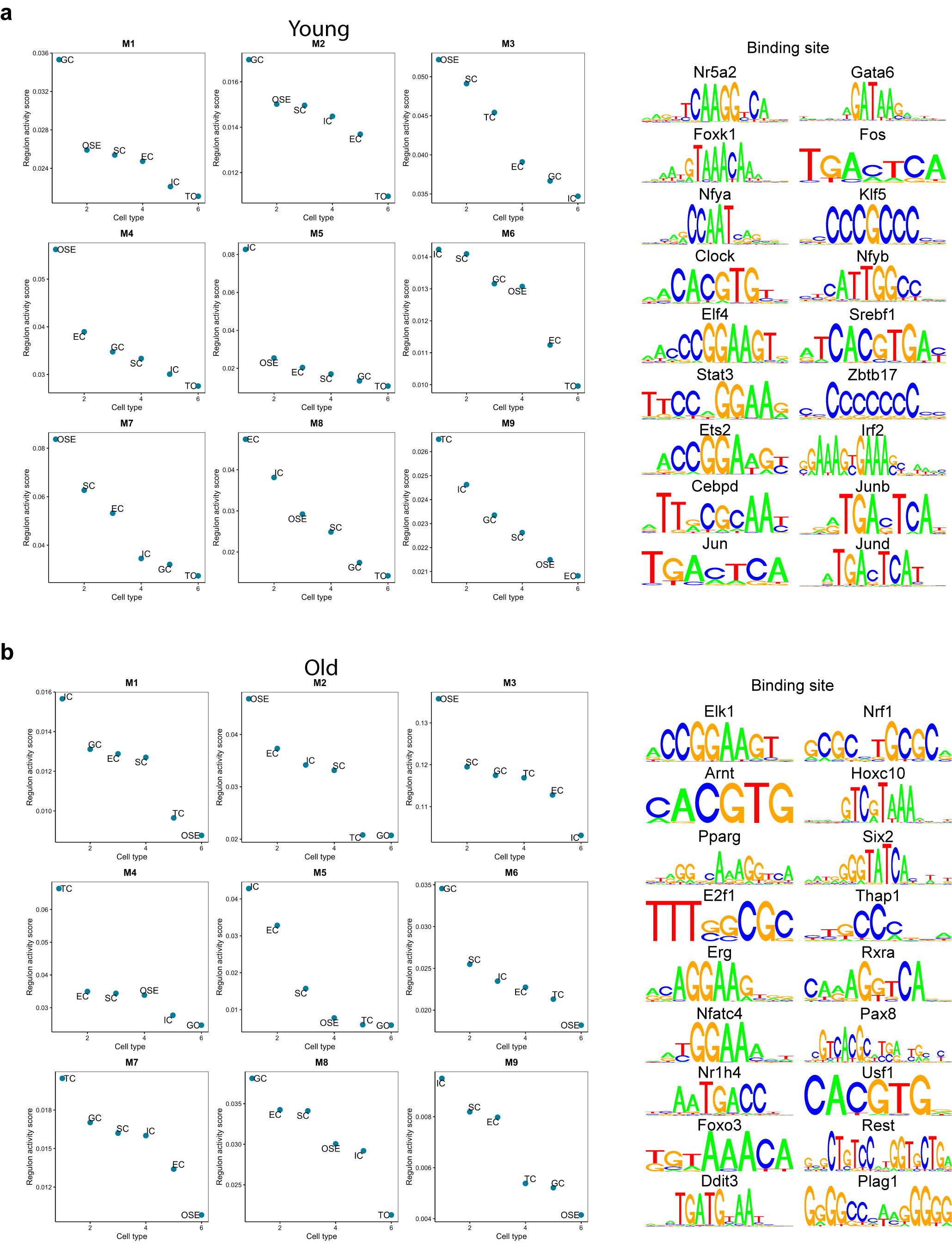

### FigS6

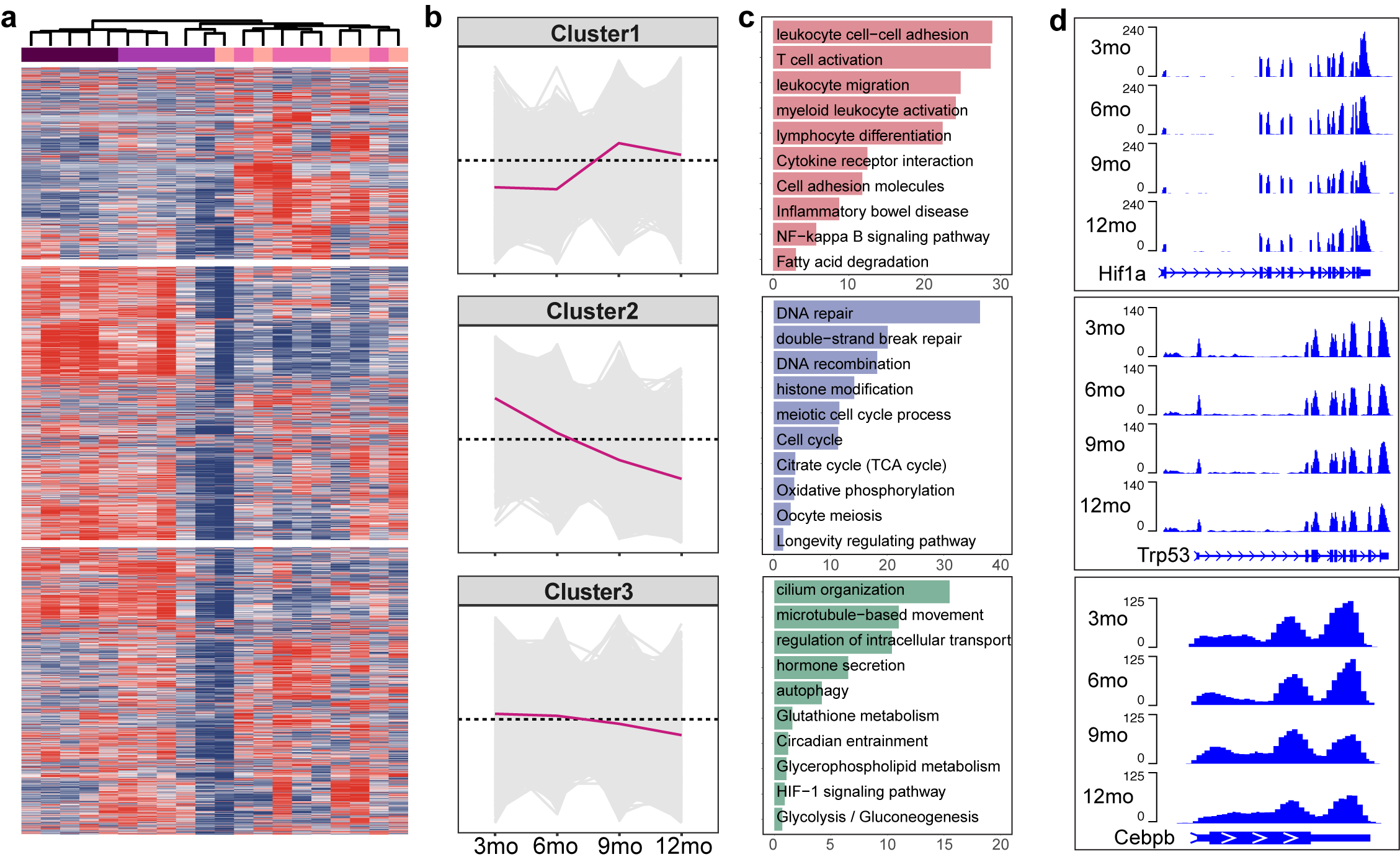

### FigS7

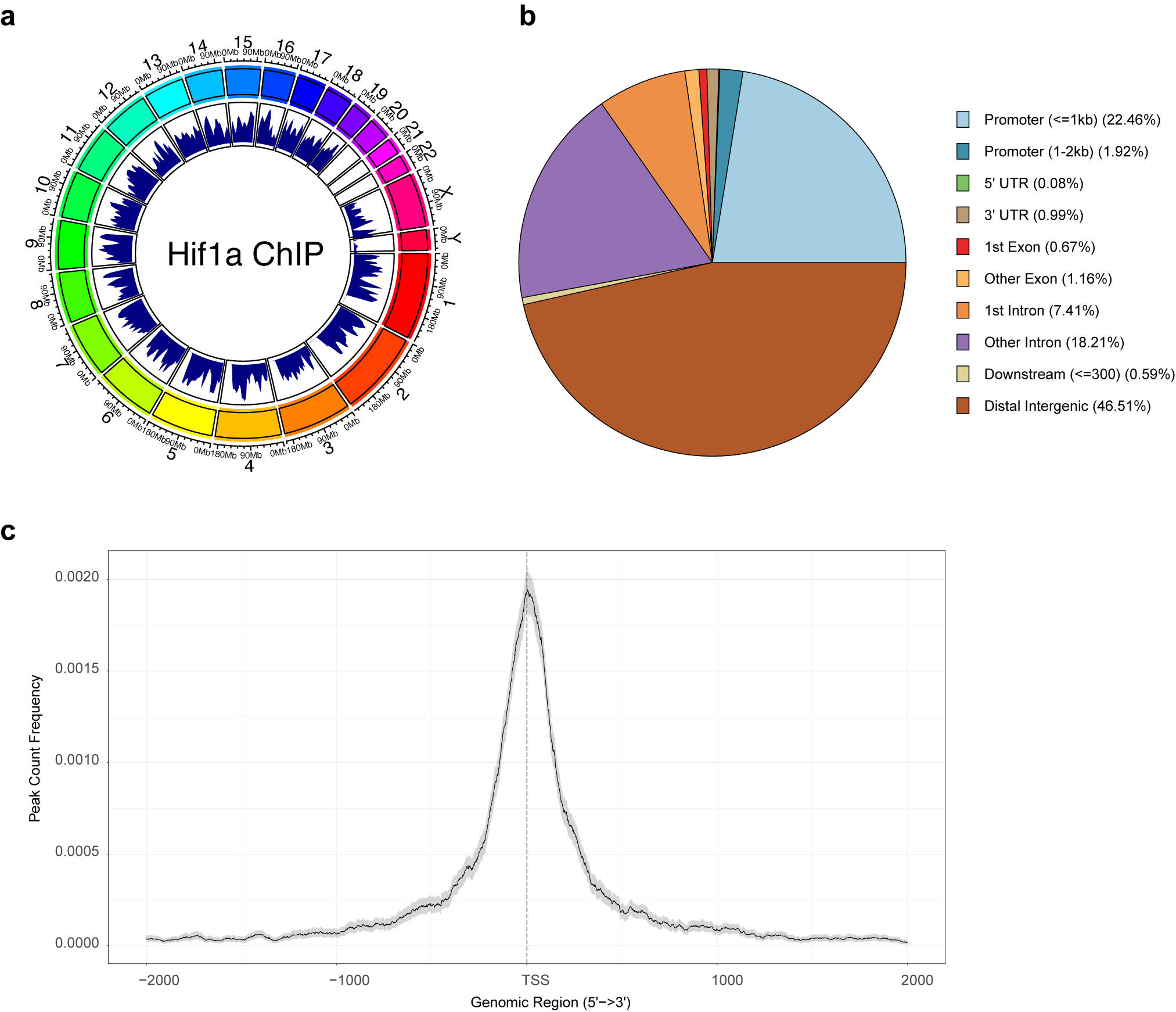
